# Hierarchical tissue structure creates history-dependent barriers to clonal invasion

**DOI:** 10.64898/2026.08.04.742588

**Authors:** Timmy Ma, Angela G. Fleischman, Dominik Wodarz, Natalia L. Komarova

**Affiliations:** Department of Mathematics, Xavier University of Louisiana, New Orleans, LA; Department of Medicine, University of California Irvine, Irvine, CA; Department of Ecology, Behavior and Evolution, University of California San Diego, La Jolla, CA; Department of Mathematics, University of California San Diego, La Jolla, CA

## Abstract

Tissues of higher organisms are maintained by hierarchies of stem and progenitor cell compartments regulated by homeostatic feedback. Somatic mutations generate genetically distinct clones whose evolutionary success depends not only on their fitness but also on the tissue architecture in which they arise. In previous work, we showed that this hierarchical organization creates invasion barriers that prevent advantageous mutants originating in downstream compartments from expanding unless their fitness exceeds a critical threshold. Here, we extend this framework to populations containing multiple competing mutant clones. We derive a general invasion criterion showing that the threshold for mutant expansion is determined by the equilibrium established by the resident clones and therefore depends on the evolutionary history of the system. Established clones modify the invasion barriers encountered by subsequent mutants, making clonal evolution history-dependent. The theory predicts competitive exclusion between clones entering the same compartment and shows that resident clones can prevent the establishment of later mutants. Using a model previously parameterized for murine hematopoiesis, we showed that our framework provides a mechanistic explanation for mutation-order effects involving JAK2 V617F and TET2 mutations in myeloproliferative neoplasms. Our results identify invasion barriers as a principle governing history-dependent clonal evolution in hierarchical tissues.

## 1 Introduction

Many adult tissues are maintained by a hierarchical organization of stem and progenitor cells, in which a small population of long-lived, multipotent stem cells gives rise to increasingly differentiated progenitors that ultimately produce the mature, functional cells of the tissue. Cell-fate decisions within such hierarchies are regulated by homeostatic mechanisms that balance self-renewal and differentiation, thereby maintaining approximately stable tissue output throughout life. At the same time, because tissue maintenance depends on the continual division of stem and progenitor cells, these tissues are inherently evolutionary systems: somatic mutations accumulate over time, and mutations that confer a competitive advantage can drive the clonal expansion of a mutant cell and its descendants at the expense of their neighbors. As a consequence, the clonal composition of a tissue is not fixed, but can change substantially as an organism ages.

This phenomenon has been documented directly through deep genomic sequencing of morphologically normal tissue. Positively selected clones carrying mutations in known cancer-driver genes progressively colonize sun-exposed skin [1], the esophageal epithelium [2, 3], and the colorectal epithelium [4], among other tissues, often occupying a substantial fraction of the tissue by middle or old age without any accompanying histological evidence of malignancy. These studies establish that age-related clonal evolution driven by somatic selection is a general feature of hierarchically structured tissues, rather than a phenomenon confined to cancer.

Among these tissues, hematopoiesis has been studied in the greatest detail, owing to the accessibility of blood and bone marrow for sampling and to the comparatively well-characterized hierarchy of stem and progenitor compartments that sustains blood-cell production, ranging from long-term hematopoietic stem cells (LT-HSCs) to increasingly differentiated progenitors. In this system, age-related clonal expansion of mutant hematopoietic stem or progenitor cells is referred to as clonal hematopoiesis: the disproportionate contribution of a single hematopoietic stem or progenitor cell and its descendants to the circulating blood-cell population. Its prevalence increases with age, and it is frequently associated with somatic mutations that alter the fitness of hematopoietic stem cells [5, 6]. The term clonal hematopoiesis of indeterminate potential (CHIP) is used when a hematologic malignancy-associated mutation is detected above a specified clone-size threshold in an individual who does not meet diagnostic criteria for a hematologic neoplasm or another defined clonal blood disorder [7]. Mutations in epigenetic regulators such as DNMT3A, TET2, and ASXL1 account for a large proportion of CHIP. Although CHIP is not itself a malignancy, it is associated with an increased risk of hematologic cancer [5, 6], cardiovascular disease [8], and other inflammatory disorders.

Mathematical models have long played an important role in understanding hematopoietic stem cell dynamics, homeostatic regulation, and clonal evolution. Classical compartmental and feedback-regulated descriptions of hematopoiesis were developed [9, 10, 11, 12], and further established a broader mathematical framework for normal and malignant hematopoiesis, including stem-cell self-renewal, leukemic progression, and competition between cell populations [13, 14, 15, 16, 17, 18, 19, 20, 21]. Evolutionary approaches have focused on the accumulation and selection of somatic mutations in hematopoietic tissues [22, 23, 24, 25, 26, 27, 28]; see also [29] for a recent perspective on the field.

A parallel body of quantitative work has modeled clonal dynamics in other self-renewing tissues, largely independently of this hematopoietic modeling tradition. A general framework for stochastic stem-cell fate in cycling tissues was developed to interpret lineage-tracing data [30], and was used to demonstrate neutral-drift dynamics of stem-cell replacement in the intestinal crypt [31]; a related population-genetics framework has been used to infer rates of stem-cell self-renewal and the strength of positive selection acting on mutant clones directly from deep-sequencing data in human epidermis [32]. These models typically treat the tissue as a single, unstructured progenitor compartment, rather than the multi-compartment, feedback-regulated hierarchy that has been central to hematopoietic modeling.

In a recent study, we developed an experimentally parameterized mathematical model of hematopoiesis that incorporated homeostatic feedback regulation and clonal evolution [33]. We showed that the hierarchical structure of hematopoiesis fundamentally changes the conditions for mutant invasion. In particular, mutants arising in downstream compartments face invasion barriers: even if they possess a fitness advantage, they cannot invade unless that advantage exceeds a critical threshold. These barriers emerge because resident populations benefit from a continuous influx of cells from upstream compartments, whereas newly arising mutants do not. As a consequence, the hematopoietic system exhibits a significant degree of resilience to mutant invasion.

The previous analysis focused on the invasion of a single mutant clone into an otherwise wild-type population. In reality, however, tissue evolution takes place in the presence of multiple competing lineages, whose fitness advantages, compartments of origin, and mutual ecological interactions can all differ. The expansion of one clone can alter the environment experienced by another. The existence of several mutant clones therefore raises new questions that a single-mutant analysis cannot address: How do invasion barriers change when other mutants are already present? Does the success of a clone depend only on its own fitness advantage, or also on the evolutionary history of the system? Can an early-arriving mutant prevent the establishment of a fitter clone that appears later, or conversely, can a newly arising clone eliminate previously established mutants? Understanding how such interactions, and, in particular, the order in which mutations are acquired, shape evolutionary outcomes has therefore become an important problem in the study of clonal evolution.

In this paper, we extend the invasion-barrier framework of [33] to the dynamics of multiple advantageous mutant clones within a hierarchically structured tissue. As a case-study, we use our model calibrated with experimental data from mouse hematopoiesis [33], which provides one of the few systems in which the kinetics of a multi-compartment stem-cell hierarchy have been quantitatively characterized. The underlying model structure, however (a hierarchy of feedback-regulated compartments linked by differentiation) is not specific to blood, and the results we derive apply in principle to any tissue organized along similar lines. We consider mutants that arise in different compartments of the hierarchy and investigate how their ability to invade depends on the composition of the resident population. We show that invasion barriers are no longer determined only by the properties of a compartment. Instead, they depend on the collection of clones that are already present and, crucially, on the compartments in which these clones originated. Thus, clonal interference in hierarchically structured tissues acquires a distinctly hierarchical component: competition between clones is shaped not only by fitness differences but also by the location within the hierarchy where mutations first arise.

This framework leads to a description of hematopoietic cell evolution in which historical contingency plays a central role. The order of mutation acquisition and the compartments in which mutations originate can alter the future accessibility of evolutionary trajectories. Consequently, the success or failure of a mutant clone depends not only on its intrinsic fitness but also on who got there first. We illustrate this framework with an application to sequential mutation acquisition in hematopoiesis, showing how the order in which advantageous mutations are acquired can determine evolutionary trajectories.

## 2 Model formulation and the concept of invasion barriers

### 2.1 The basic model and parameters

We consider a mathematical model of tissue cell homeostasis that describes the dynamics of cell populations across a hierarchy of compartments, ranging from stem cells to progressively more differentiated progenitors. The model consists of a system of ordinary differential equations that track the abundances of cells in each compartment. Cell populations evolve through division events that can result either in self-renewal or differentiation, with division rates denoted by *r*_*i*_ and self-renewal probabilities given by functions *P*_*i*_(·) that depend on the cell population sizes.

Specifically, we consider four tissue compartments. *C*_0_ denotes the most upstream stem cells, and compartments *C*_1_, *C*_2_, and *C*_3_ denote progressively differentiated stem / progenitor cells. While this mathematical modeling structure can apply to a variety of hierarchically organized tissues, we will parameterize the model for murine hematopoiesis because of the availability of detailed kinetic data from label propagation experiments that allow parameter estimations of hematopoietic stem cells (HSC) dynamics [34, 33]. In this context, compartments *C*_0_ – *C*_3_ correspond to long term (LT)-HSCs, short term (ST)-HSCs, multipotent progenitors (MPPs), and the sum of commmon myeloid progenitors (CMPs) and common lymphoid progenitors (CLPs), see table 1.

**Table 1:** Hematopoietic stem cell compartments and the mathematical notations. The mathematical model allows for an arbitrary number of downstream compartments, *n*.

|  |  |  |  |  |
| --- | --- | --- | --- | --- |
| Compartment notation | $C_0$ | $C_1$ | $C_2$ | $C_3$ |
| Cell type | LT-HSCs | HT-HSCs | MPPs | CMPs and CLPs |

Denote by *x*_*i*_(*t*) the number of cells in compartment *i*, respectively. The system of equations that describes the co-dynamics of these cells is given by (see also the schematic in figure 1):

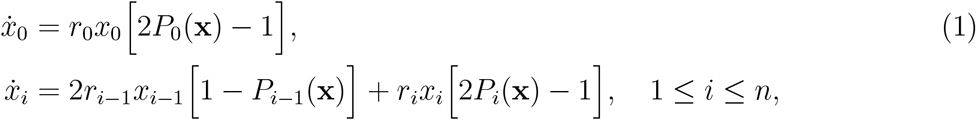

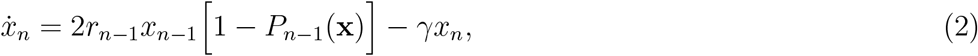

where **x** = (*x*_0_, …, *x*_*n*_)^*T*^, see [33]. A key feature of the model is the presence of feedback control, whereby the effective self-renewal probability decreases as the population size within a compartment increases. This negative feedback stabilizes the system and gives rise to a homeostatic equilibrium in which all compartments are maintained at steady levels [33].

**Figure 1:**
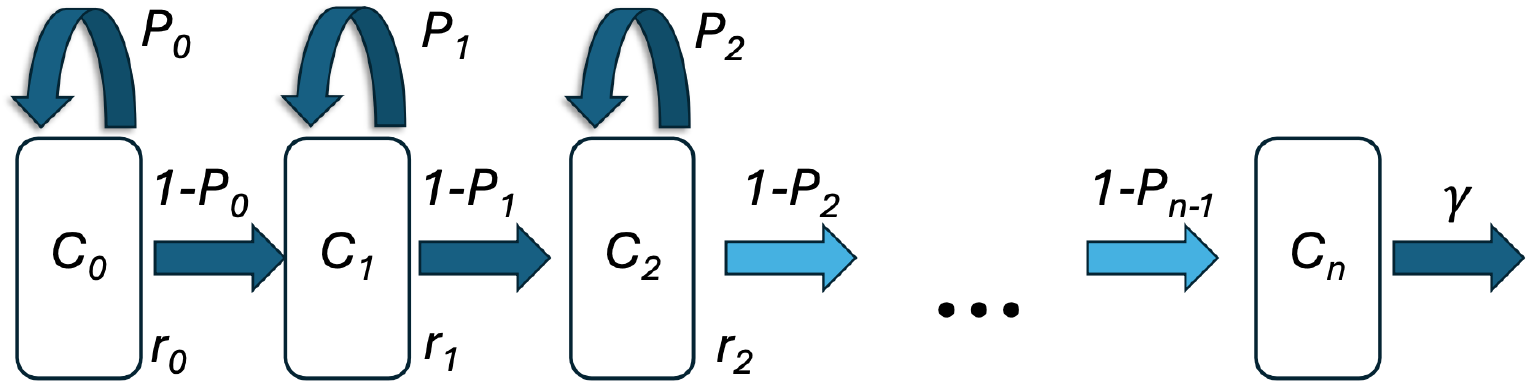
A schematic showing a hierarchical organization of the hematopoietic system used in the model.

While the functions *P* (**w**) can be of many different forms [13], following [33], for our numerical illustrations here we will consider a special case of the dependencies of functions *P*_*i*_ on cell populations. Namely, we will assume that control comes from within each compartment:

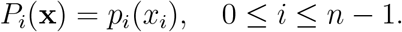

In particular, we will use the following functional form:

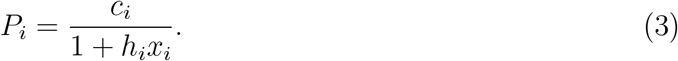

Using the experimental data reported in [34], we previously estimated kinetic parameters of system (1-2) at steady state, see [33]. These parameters include the division rates of cells in the different compartments, *r*_*i*_, and the *equilibrium* self-renewal rates, 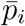, see table 2. These are the values of the functions *P*_*i*_(**x**) evaluated at the equilibrium; importantly, the equilibrium values reflect the experimental data and do not depend on our assumptions on the functional form of *P*_*i*_(**x**). The equilibrium population sizes are given by

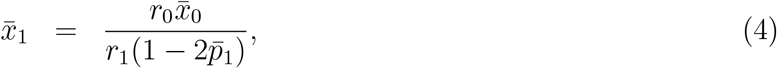

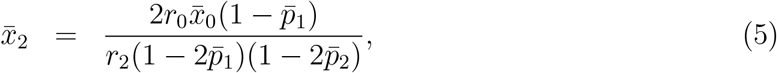

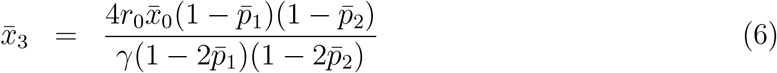

**Table 2:** Definitions of parameters and their values, taken from [33] and based on the experiments in [34].

| Notation | Parameter definition | Value | 95% C.I. | Units |
| --- | --- | --- | --- | --- |
| $r_0$ | Division rate of LT-HSCs | 0.0107 | (0.0084, 0.014) | days<sup>-1</sup> |
| $r_1$ | Division rate of ST-HSCs | 0.067 | (0.021, 0.10) | days<sup>-1</sup> |
| $r_2$ | Division rate of MPPs | 0.136 | (0.066, 0.95) | days<sup>-1</sup> |
| $\bar{p}_0$ | Self-renewal probability of LT-HSCs | 0.5 | | 1 |
| $\bar{p}_1$ | Self-renewal probability of ST-HSCs | 0.473 | (0.40, 0.48) | 1 |
| $\bar{p}_2$ | Self-renewal probability of MPPs | 0.416 | (0.323, 0.49) | 1 |
| $\gamma$ | Removal rate of CMPs+CLPs | 0.0274 | (0.015, 0.17) | days<sup>-1</sup> |
| $\bar{x}_0$ | Equilibrium number of LT-HSCs | 17,000 | | cells |

Using a specific assumption about the functional form of the self-renewal probabilities, equation (3), we have the following relationship between parameters *h*_*i*_ and *c*_*i*_:

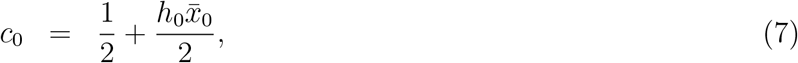

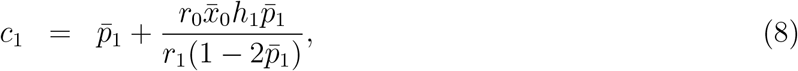

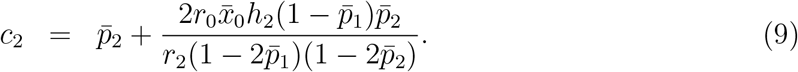

### 2.2 Multiple clones and system extensions

System (1-2) can be adapted to describe co-dynamics of multiple clones that inhabit hematopoietic compartments. For example, assuming that *x*(*t*) describes the size of the wild-type population and *y*(*t*) the size of the mutant population, we can write

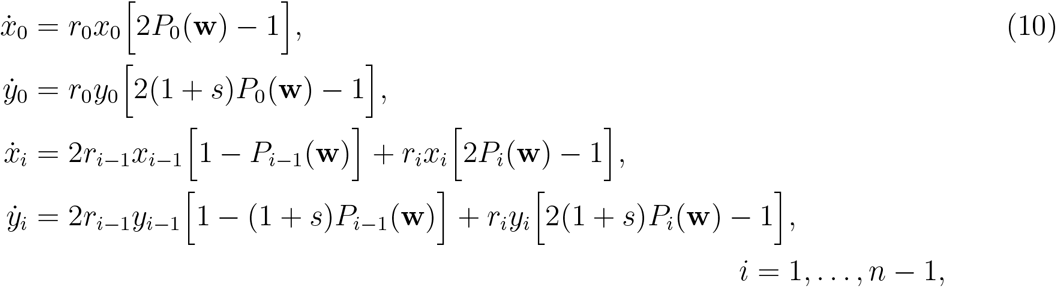

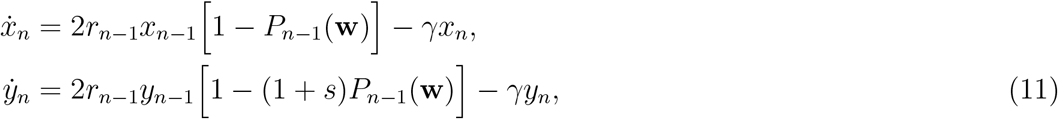

where *w*_*i*_ = *x*_*i*_ + *y*_*i*_ and **w** = (*w*_0_, …, *w*_*n*_)^*T*^, see [33]. In this model the different clones differ from each other in their effective self-renewal probability. In particular, if *y*(*t*) is an advantageous mutant, then it is assumed that it has an increased tendency to self-renew, expressed as a multiplicative factor (1 + *s*) applied to the self-renewal probability. Importantly, both clones experience the same feedback control, so that they compete for the same regulatory environment.

A generalization to any number of clones is straightforward. Let us suppose there are *K* different mutant clones, each clone *k* with its selection coefficient *s*_*k*_ ≥ 0, and use the superscripts to enumerate the different clones. We have:

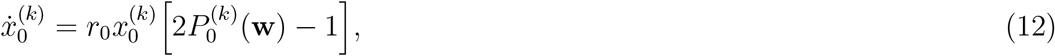

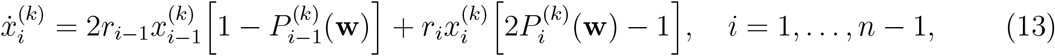

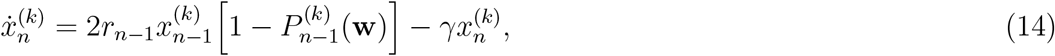

where 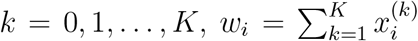, and 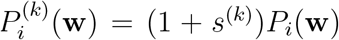 is the rate of self-renewal experienced by clone *k* in compartment *i*. We will assume that clone *k* = 0 is the wild-type, and set *s*^(0)^ = 0. The rest of the clones measure their selection coefficient *s*^(*k*)^ relative to the wild-type clone, *k* = 0.

The question of interest is whether a mutant population of a given type introduced at low numbers can grow and establish itself within the system. Ultimately, the answer to this question depends on (1) what compartment is the new clone introduced into and (2) what other clones are already there.

In system (12-14) there is no explicit *de novo* production of mutants. Instead, we will introduce new clones by means of initial conditions. For example, we would use the following initial conditions to model the situation where a single mutant of type *k* = 1 is produced in compartment *C*_*i*_ with *i* = 2 at time *t* = *t*_0_:

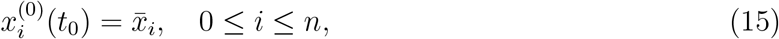

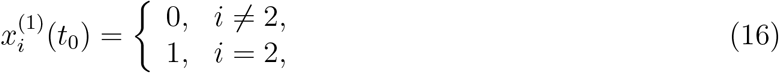

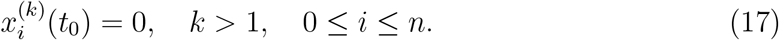

For a model that explicitly includes mutant production, please see [33].

### 2.3 Invasion barriers: the first mutant clone

Previously we considered scenarios where a single mutant is introduced in a given compartment of a system that is at a homeostatic equilibrium [33]. We demonstrated that the presence of hierarchical compartment structure fundamentally alters the conditions for mutant invasion. In particular, the system exhibits a form of resilience to invasion, characterized by the existence of invasion barriers. These barriers arise when mutants originate in any of the downstream compartments, *C*_*i*_ with *i >* 0, such as short-term stem cells or multipotent progenitors. In such cases, mutant cells lack the continuous influx from upstream compartments that contributes to the effective fitness of the resident population. As a result, even if mutants possess a higher intrinsic self-renewal probability, they may still fail to invade unless their advantage exceeds a critical threshold.

This threshold can be understood by considering the effective growth rate of a rare mutant in a given compartment. For example, if a mutant arises in one of the downstream compartments, *C*_*k*_ with *k >* 0, invasion requires a condition of the form

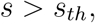

where the threshold *s*_th_ depends on the equilibrium value of the self-renewal probability.

For system (10-11) that describes the co-dynamics of wild-type cells and mutants with the selection coefficient *s*, the invasion barrier condition for a mutant that is first introduced in compartjment *C*_*i*_ is given by

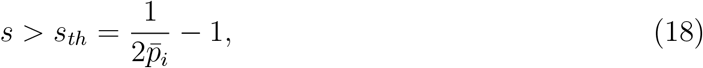

where 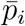 is the equilibrium self-renewal probability of the wild-type cells, see table 2. In the parameter regime relevant to hematopoiesis (in the mouse system), these thresholds can be substantial. In particular, mutants arising in the ST-HSC compartment (*C*_1_) require fitness advantages higher than 5.7% (i.e., *s*_*th*_ ≈0.057), and for the MPP (*C*_2_) we have *s*_*th*_ ≈0.202, such that an advantage of more than 20.2% is required to invade. Mutants with smaller advantages will be eliminated.

By contrast, mutants originating in the most upstream compartment do not experience such barriers. In that case, invasion is determined simply by a comparison of fitness parameters: the type with the larger effective self-renewal probability ultimately dominates the population. This can be seen from equation (18), which for *i* = 0 gives *s*_*th*_ = 0 because 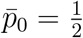. Thus, the system exhibits a strong asymmetry with respect to the location of mutant origin.

At the same time we note that the likelihood of generating a mutation is the lowest for the upstream compartments (because of a much lower compartment size), and if a mutant originates e.g. in the top compartment (LT-HSC, *C*_0_), it will rise extremely slowly due to a slower division rate there.

The existence of invasion barriers can be interpreted as a consequence of the interplay between feedback control and hierarchical organization. The influx of cells from upstream compartments effectively enhances the persistence of the resident population in downstream compartments, thereby creating a disadvantage for newly arising mutants.

### 2.4 Equilibrium solution types

Suppose all possible clones are enumerated, with the wild-type assigned index 0. Multiple clones can coexist at equilibrium in system (12-14), and the equilibrium depends both on the selection coefficients *s*^(*k*)^ ≥ 0 and on the compartments in which the different clones first arose. Here we introduce a simple notation that summarizes the resulting clonal architecture.

If an advantageous mutant is introduced into the top compartment, *C*_0_, it eventually displaces the wild-type and becomes the new background clone, see figure 2(a). If instead a mutant is introduced into a downstream compartment *C*_*i*_ with *i >* 0, and its selection coefficient is above the corresponding invasion barrier, it establishes itself in *C*_*i*_ and all downstream compartments, but never in compartments above *C*_*i*_, see figure 2(b,c). Thus, if clone *k* first establishes in compartment *C*_*i*_, we have

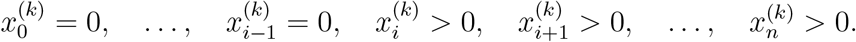

**Figure 2:**
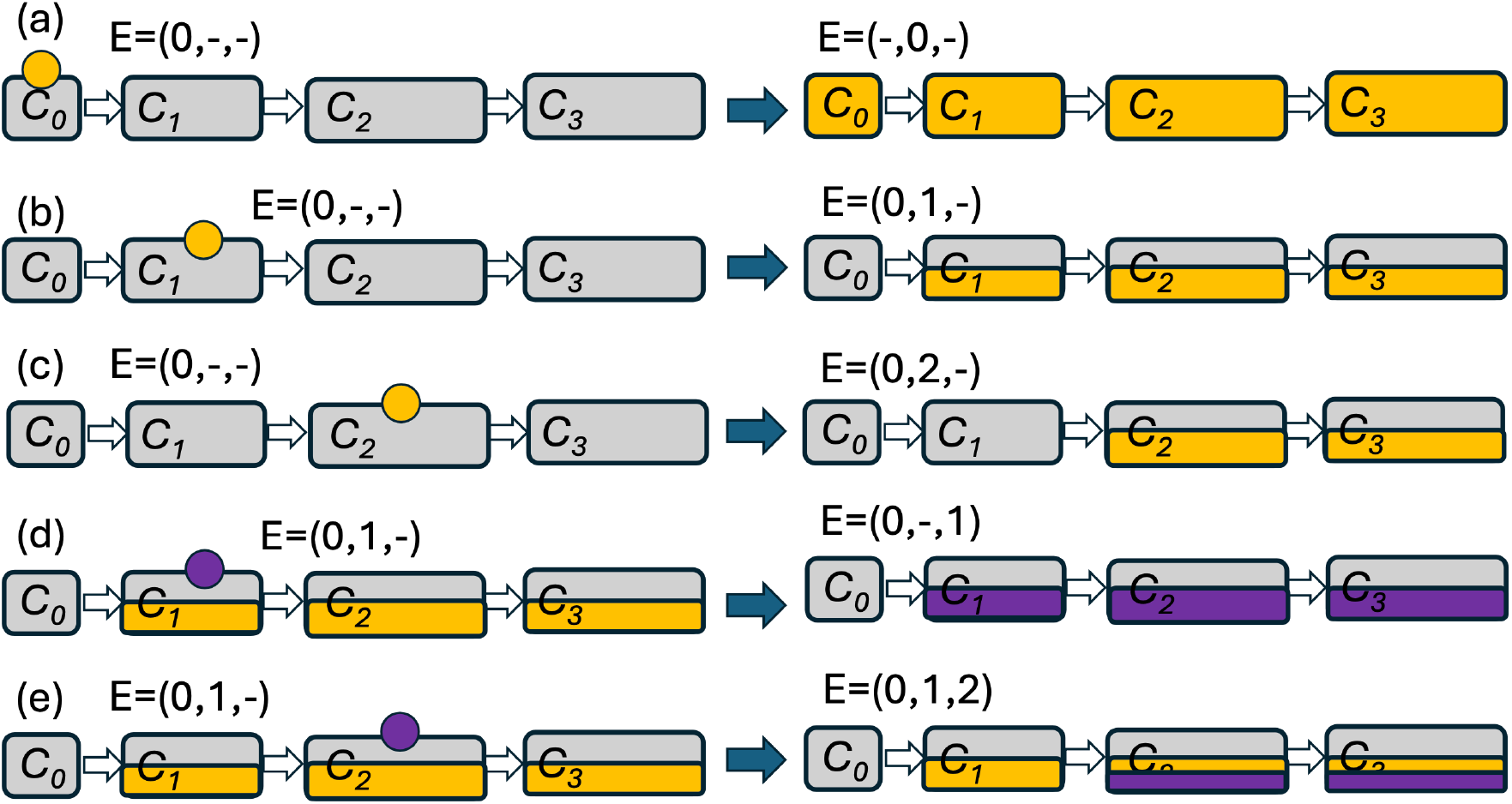
A schematic showing different types of clonal dynamic. Each panel represents an initial condition on the left and the resulting equilibrium on the right. The gray color corresponds to the wild-type clone (*k* = 0), the yellow and purple are clones with *k* = 1 and *k* = 2 respectively. A circle above a compartment shows that at the initial time, a small amount of the corresponding type was introduced in that compartment. It is assumed that the new clone’s selection coefficient is always above the relevant invasion barrier (otherwise the new mutant would go extinct). Multiple colors within a compartment indicate coexistence of different types. In panels (a,b,c), initially a small amount of clone *k* = 1 was introduced in compartments *C*_0_, *C*_1_, *C*_2_ respectively. In panels (d,e), clone *k* = 2 was introduced into compartments *C*_1_ and *C*_2_, respectively, while clone *k* = 1 was already residing there. The entry strings are shows for each resident equilibrium on the left and for each resulting equilibrium on the right, before and after the new clone was introduced.

This motivates the following definition.

We define the *entry compartment* of clone *k* as the first compartment in which that clone is present,

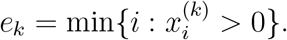

Equivalently,

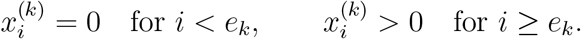

Because cells differentiate only downstream, the entry compartment uniquely determines the qualitative compartmental distribution of a clone within the hierarchy.

For example, in figure 2, the gray clone (wild-type) has index *k* = 0, the yellow clone has index *k* = 1, and the purple clone has index *k* = 2. In panel (b), the yellow clone first appears in compartment *C*_1_, so *e*_1_ = 1. In panel (c), it first appears in compartment *C*_2_, so *e*_1_ = 2. Throughout figure 2, successful invasion is assumed.

We summarize an equilibrium by its *entry string*,

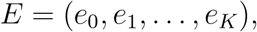

which simply lists the entry compartment of every clone. If clone *j* is absent from the system, we write *e*_*j*_ = −. Thus, the entry string provides a compact description of the equilibrium clonal architecture.

For example,

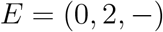

means that the wild-type occupies the system from *C*_0_, the first mutant (*k* = 1) entered in compartment *C*_2_, and the second mutant (*k* = 2) is absent, see figure 2(c). Likewise,

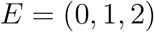

(figure 2(e)) indicates that mutant *k* = 1 entered in compartment *C*_1_ and mutant *k* = 2 entered in compartment *C*_2_. The position of each entry in the string identifies the clone, while its value specifies the compartment where that clone first became established.

### 2.5 A general case: invasion barriers depend on who got there first

Here we generalize the results of [33] (section 2.3) to the existence of multiple advantageous clones. Consider system (12-14). Suppose that the wild-type and possibly some mutant clones coexist non-trivially in the system, and their entry compartments are given by a string

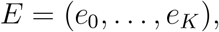

which has both nontrivial and “−” entries. We assume that the clone of interest, which we call clone *ℓ*, is not yet in the system, *e*_*ℓ*_ = −. Denote by 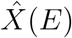 this (resident) equilibrium as 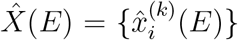 with 0 ≤ *i* ≤ *n*, 0 ≤ *k* ≤ *K*. The mutant clone (*k* = *ℓ*) is introduced in compartment *i*. Will it be able to invade in this model?

Examples of such scenarios are in figure 2(d,e): a new clone (purple) is introduced on the background of the yellow clone coexisting with the wild-type. In the schematic we show successful invasion of the purple clone. In this section we discuss the conditions for such successful invasion.

Let us denote by ***ŵ*** (*E*) the vector of total populations of the coexisting clones: ***ŵ*** (*E*) = (*ŵ*_1_(*E*), … *ŵ*_*n*_(*E*)) with 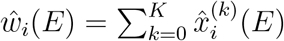. The following theorem generalizes the invasion criterion derived in [33] from a single mutant to an arbitrary resident clonal architecture.

#### Theorem 1.

For clone *ℓ* to be able to invade in compartment *i*, the following threshold condition must hold:

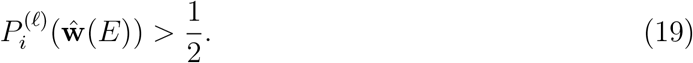

In other words, the new clone must be self-sustaining in compartment *i*, under the self-renewal rate corresponding to the resident equilibrium.

**Proof**. To derive condition (19), suppose that the resident clones are at the equilibrium 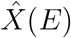, and introduce clone *ℓ* at low abundance in compartment *C*_*i*_, with no cells of this clone in upstream compartments: 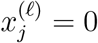 for *j < i*. Because differentiation is unidirectional, these upstream populations remain zero. The equation for the new clone in its entry compartment is therefore

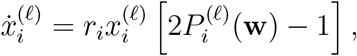

since the influx term from compartment *C*_*i*−1_ vanishes.

When the invading clone is rare, it has a negligible effect on the feedback environment, so that

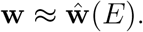

Linearizing around the resident equilibrium gives

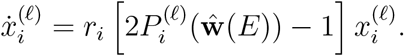

Thus, the per-capita growth rate of the rare clone in its entry compartment is

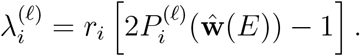

Since *r*_*i*_ *>* 0, the clone grows from low abundance in compartment *C*_*i*_ if and only if

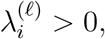

or equivalently,

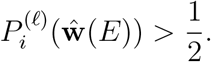

If the inequality is reversed, the clone cannot expand from low numbers in the entry compartment. □

As the first example, consider the case where the resident population is the wild-type, the entry string is just *E* = *E*_0_ ≡ *{*0, −, …, −*}*, and the clone that we introduce is *ℓ* = 1. For *i* = 0 (figure 2(a)), the self-renewal probability of the wild-type satisfies 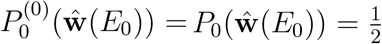 and therefore inequality (19) requires

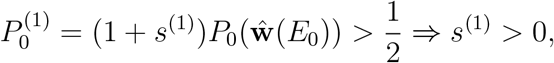

that is, any advantageous mutant can invade from compartment *C*_0_. For the downstream compartments, 1 *< i < n* (figure 2(b,c)), we observe that 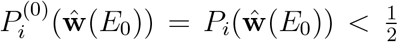 (this follows from equation (13) at the equilibrium) and thus

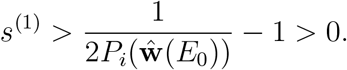

This is exactly the result obtained in [33] that shows the existence of a non-trivial invasion barrier in downstream compartments. In particular, the explicit expression for the invasion barrier in compartment *C*_1_ under our specific functional form of self-renewal probability, is given by

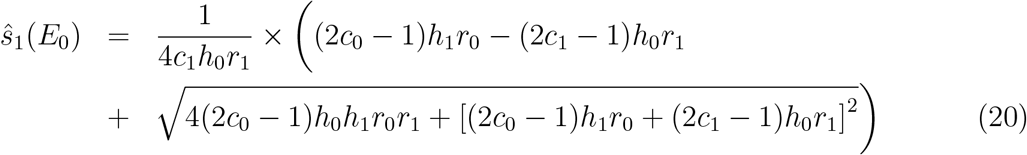

An explicit expression for *ŝ*_2_(*E*_0_) can also be obtained, but it is longer.

Next, consider the case where more than one resident clones are present. Condition (19) gives

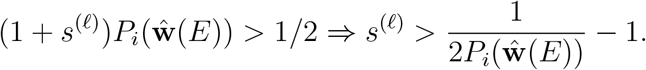

Let us denote by *ŝ*_*i*_(*E*) the threshold value for the selection coefficient of a mutant clone that makes invasion from low numbers possible, given that it enters an established equilibrium characterized by the entry string *E*. We have

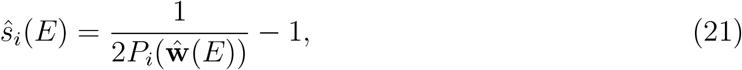

and clone *ℓ* can expand from low numbers in compartment *i* if *s*^(*ℓ*)^ *> ŝ*_*i*_(*E*). We can distinguish the following two scenarios:

1. Compartment *i* is an entry compartment for one of the resident clones, *m*: *e*_*m*_ = *i*, see e.g. figure 2(d), where clone *ℓ* = 2 enters compartment *C*_1_, which is the entry compartment for the resident clone *m* = 1: *e*_1_ = 1;
2. Compartment *i* is not an entry compartment for any of the resident clones, that is, *e*_*k*_ ≤ *i* for all *k*, see e.g. figure 2(e), where clone *ℓ* = 2 enters compartment *C*_2_, which is not an entry compartment for any of the resident clones (*e*_0_ = 0, *e*_1_ = 1).

Below we denote the newly introduced clone as *ℓ*.

#### 1. A new clone *ℓ* is introduced in an entry compartment of another clone, *m*

For clone *m*, 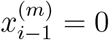, and we have in system (13) at the equilibrium, in *C*_*i*_,

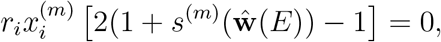

which means that 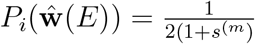. Substituting this into equation (21), we get

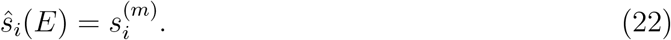

This means that the new clone that is introduced into compartment *i* must have a selection coefficient that is larger than that of the resident clone that entered in that compartment.

Once a new clone successfully enters the system in a compartment that was an entry compartment for one of the resident clones, this will result in a competitive exclusion. This follows from the following simple argument. Suppose a resident clone *m* had entered in compartment *C*_*i*_, and then clone *ℓ* also entered in *C*_*i*_ and expanded successfully. This means that

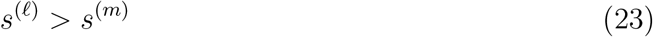

because the threshold (22) must have been overcome. Suppose these two clones coexist in a stable equilibrium. Then from system (13) it follows that

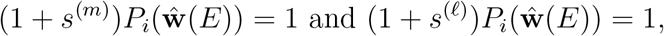

which is impossible because *s*^(*ℓ*)^ *> s*^(*m*)^. Therefore the stronger clone will displace the weaker clone leading to competitive exclusion.

Furthermore, if the new clone *ℓ* displaced *m*, it will spill over to the downstream clones, where it will also displace clone *m*, leading to its eventual extinction in the whole system. For example, consider compartment *C*_*i*+1_, immediately downstream from the compartment where the new clone *ℓ* displaced the resident clone *m*. We have from system (13),

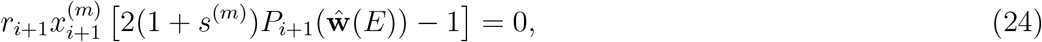

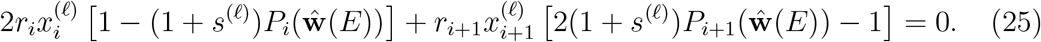

From equation (24) we obtain, if 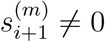,

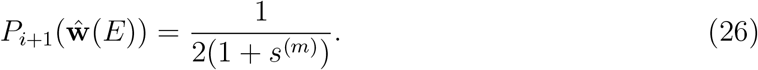

From equation (25), since we assumed 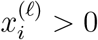, we have

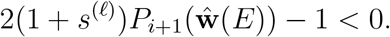

Using (26) we obtain *s*^(*ℓ*)^ *< s*^(*m*)^, which is a contradiction. Therefore, we must have *x*^(*m*)^ = 0. In other words, the winning clone *ℓ* that excluded clone *m* in its entry compartment will eliminate it in the entire system. Figure 2(d) illustrates that: the purple clone (assumed stronger than the yellow clone) enters in the same compartment *C*_1_ and eliminates the yellow clone in *C*_1_ and everywhere downstream.

Finally, the new clone *ℓ* might displace additional clones with entry compartments down-stream from *C*_*i*_. Let us denote the original entry string (for the state with clone *m*) as *E*_1_, and the new entry string where *ℓ* displaced *m* as *E*_2_. Then a clone *q* with *e*_*q*_ = *j > i* will be displaced if (1 + *s*^(*q*)^)*P*_*j*_(*E*_2_) *<* 1*/*2. This can happen, for example, if the population size in *C*_*j*_ increased after the displacement of *m* with a fitter *ℓ*, enough to lower *P*_*j*_(*E*_2_) below 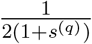.

#### A new clone *ℓ* is introduced in a compartment that is not an entry compartment of any other clone

Let us suppose that *k* is a clone with an entry compartment upstream from *i*, and the new clone *ℓ* enters in *C*_*i*_. Then from system (13) for compartment *i* we can see that

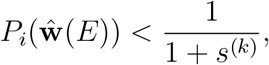

and therefore, using expression (21), we obtain that

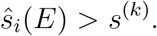

In other words, the new clone in compartment *C*_*i*_ has to overcome a barrier, which is higher than the selective coefficient of all the clones that entered upstream from *C*_*i*_.

A special case of scenario 2 is when the first mutant, *m* = 1, with *s*^(1)^ *>* 0, enters the top compartment, *C*_0_. It will invade and displace the wild-type. Mathematically it can be treated as a new “wild type”. All other clones will experience “updated” (and higher) invasion barriers, which are higher than the selection coefficient of the mutant residing in *C*_0_, *s*^(1)^. Denoting by *E*_*new*_ ≡ *{*−, 0, − …, −*}* the entry configuration that corresponds to the equilibrium with mutant *k*_1_ fixated in the system, we obtain

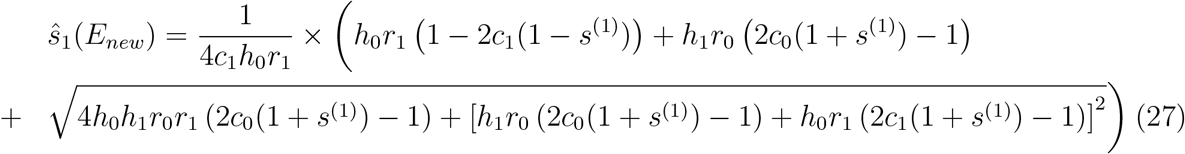

This expression replaces the barrier that corresponds to the wild type-based equilibrium, equation (20). Figure 3(a) shows the quantity *ŝ*_1_(*E*_*new*_) as a function of the resident mutant’s selection coefficient, *s*^(1)^, and compares it with *ŝ*_1_(*E*_0_) (equation (20), the threshold on the wild-type background). We can see that *ŝ*_1_(*E*_*new*_) *> s*^(1)^, and the inset shows that the difference increases with *s*^(1)^.

**Figure 3:**
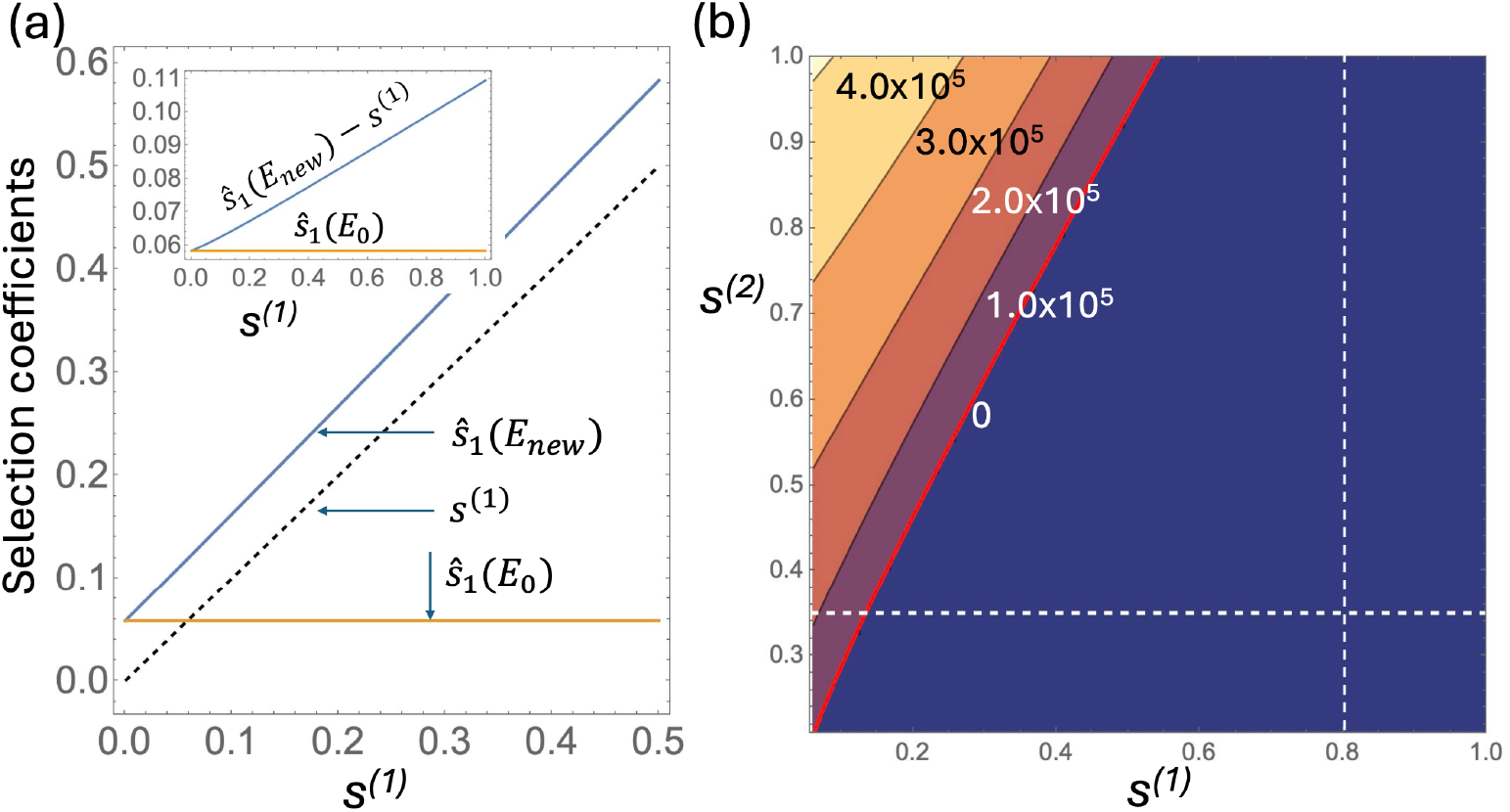
Invasion barriers. (a) The threshold value for *C*_1_ on the mutant background, *ŝ*_1_(*E*_*new*_) (equation (27), blue line), as a function of the resident clone’s selection coefficient *s*^(1)^. The threshold corresponding to the wild-type background, *ŝ*_1_(*E*_0_) (equation (20)) is shown by the yellow line and the resident clone’s *s*^(1)^ as a dashed black line. The inset shows *ŝ*_1_(*E*_*new*_) *> s*^(1)^ as a function of *s*^(1)^ in blue and *ŝ*_1_(*E*_0_) in yellow. (b) A heat plot of the equilibrium solution 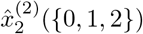, that is, the population of JAK2V617F that originates in *C*_2_, in the presence of TET2 that originated in *C*_1_. The two axes are the selection coefficient for TET2 (*s*^(1)^) and that for JAK2V617F (*s*^(2)^). Several levels are shown by contours; the zero contour (the red line) corresponds to the threshold *ŝ*_2_({0, 1, 2 }) as a function of *s*^(1)^. The left bottom corner of the graph corresponds to the point *s*^(1)^ ≈0.058 and *s*^(2)^ ≈0.21, the threshold values for the two compartments on the wild-type background. The white dashed lines mark *s*^(1)^ = 0.8 and *s*^(2)^ = 0.35. The rest of the parameters are given in table 2.

To summarize our findings, we can say that mutant dynamics in the presence of multiple clones obey several intuitive rules:

- Any new clone entering the system in compartment *C*_*i*_ experiences an invasion threshold given by equation (21). This is equivalent to condition (19), which states that the new clone is self-sustaining in *C*_*i*_ under the equilibrium dynamics of the resident clones.
- A new clone entering a system in compartment *C*_*i*_ experiences an invasion threshold that is higher than the selection coefficient of all the resident clones that have entered in *C*_*i*_ and upstream from it.
- Two resident clones cannot share the same entry compartment. The stronger clone drives the weaker clone extinct in its entry compartment and everywhere downstream from it. This is illustrated in figure 2(d): the purple clone enters in *C*_1_ (which is the entry compartment of the resident yellow clone). The purple clone is assumed to have a higher selection coefficient compared to the resident yellow clone, and it drives it extinct in *C*_1_ and in the whole system.
- A new clone that enters successfully in compartment *C*_*i*_ may coexist with other clones that entered upstream from *C*_*i*_. This is illustrated in figure 2(e): the purple clone is assumed to have a selection coefficient higher than the invasion threshold in *C*_2_, where it enters. It establishes a nontrivial population that coexists with the yellow clone (and the wild-type) in *C*_2_ and downstream from that.

## 3 Applications

We apply our theory to ask how the presence of a relatively “harmless”, non-cancerous mutant clone can affect the emergence of another, malignancy-driving mutant. This becomes especially interesting if the non-cancerous mutant has a higher self-renewal potential (and thus a higher Darwinian fitness) than the malignancy-driving mutant.

In this section we assume that there are two mutant clones. One of them (mutation *k* = 1) has a higher selection coefficient in terms of our model but is relatively “harmless”, while the other mutation (*k* = 2) has a lower selection coefficient but is more dangerous. Data indicate that this might apply to the common CHIP mutation TET2 (*k* = 1) and the JAK2V617F mutant (*k* = 2) in the hematopoietic system. We will use this particular system to explore the more general question of how a pre-existing clonal shift towards higher cell fitness (mutation *k* = 1) in a tissue can alter the potential success of a malignancy-driving mutation (*k* = 2).

The TET2 gene encodes a methylcytosine dioxygenase that plays a central role in active DNA demethylation and epigenetic regulation. Loss-of-function mutations in TET2 impair normal DNA demethylation, enhance the competitive fitness and self-renewal of hematopoietic stem cells, and are associated with altered differentiation and increased inflammatory signaling. TET2 mutations are among the most common drivers of clonal hematopoiesis of indeterminate potential (CHIP).

hsignaling. TET2 mutations are among the most common drivers of clonal hematopoiesis of indeterminate potential (CHIP).

The JAK2 gene encodes a non-receptor tyrosine kinase that transduces signals from cytokine receptors through the JAK-STAT signaling pathway to regulate normal blood cell production. The most common mutation, JAK2V617F, causes constitutive activation of the kinase, allowing signaling to occur in the absence of normal growth factor stimulation. This allows hematopoietic stem and progenitor cells to proliferate, leading to excessive production of blood cells. JAK2V617F is strongly associated with myeloproliferative neoplasms (MPNs), a group of chronic blood cancers characterized by the overproduction of one or more types of mature blood cells [35, 36].

In this paper, we assume that

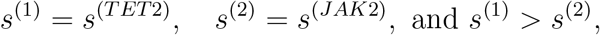

that is, the selection coefficient associated with TET2 mutations is larger than that associated with JAK2V617F mutations. In our model, the selection coefficient quantifies an increase in the effective self-renewal probability relative to the wild type, which determines competitive fitness. By contrast, an increase in proliferation rate alone does not confer a selective advantage in this framework, because self-renewal and differentiation are coupled to cell division and therefore accelerate proportionally. Experimental evidence suggests that TET2 confers the stronger self-renewal advantage. Tet2-deficient HSCs outcompete wild-type HSCs in competitive transplantation experiments and progressively expand in the stem cell compartment [37, 38]. In contrast, Jak2V617F knock-in mice develop an MPN characterized by expansion of downstream myeloid progenitors through cell division, but their mutant HSCs show no competitive advantage over wild type [39] and in some models are functionally impaired [40, 41]. Similarly, studies of patients carrying both mutations found that JAK2-first individuals overproduce mature blood cells without an accompanying self-renewal advantage at the HSC level, which appears only after acquisition of TET2, and that the HSC and progenitor-cell compartment is dominated by double-mutant rather than JAK2 single-mutant cells [42]. These observations are therefore consistent with our assumption that through higher self-renewal, the CHIP mutant TET2 might enjoy a selective advantage over a JAK2V617F mutant cell, even though the JAK2V617F mutant might be characterized by a faster rate of proliferation (which by itself does not confer a selective advantage).

We will assume that the selection coefficients of both mutants are higher than the invasion barriers in the SH-HSC and MPP compartments, that is, *s*^(*k*)^ *> ŝ*_*i*_(*{*0, −, −*}*) for *k* ∈ *{*1, 2*}* and *i* ∈ {1, 2}. In other words, if either of these mutants is placed in compartment *C*_1_ or *C*_2_ in the absence of any other mutations in the system, it will rise from low numbers. Since *ŝ*_1_({0, −,− }) ≈ 0.058 and *ŝ*_2_({0, −, −}) ≈ 0.21, we must have *s*^(1)^, *s*^(2)^ *>* 0.21, or, using biological notation,

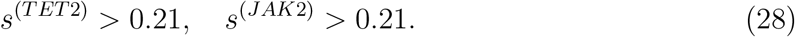

The mathematical formalism includes two separate mutations. In reality they can represent two distinct scenarios:

- The two mutational hits both arise in (two different) wild-type cells, first one and then the other, see a schematic in figure 4(a).
- The first mutation happens in a wild-type cell, spreads through the population (partially, or even reaching fixation), and then the second mutation happens in one of the mutated cells, giving rise to a double-hit mutant clone, see figure 4(b,c).

**Figure 4:**
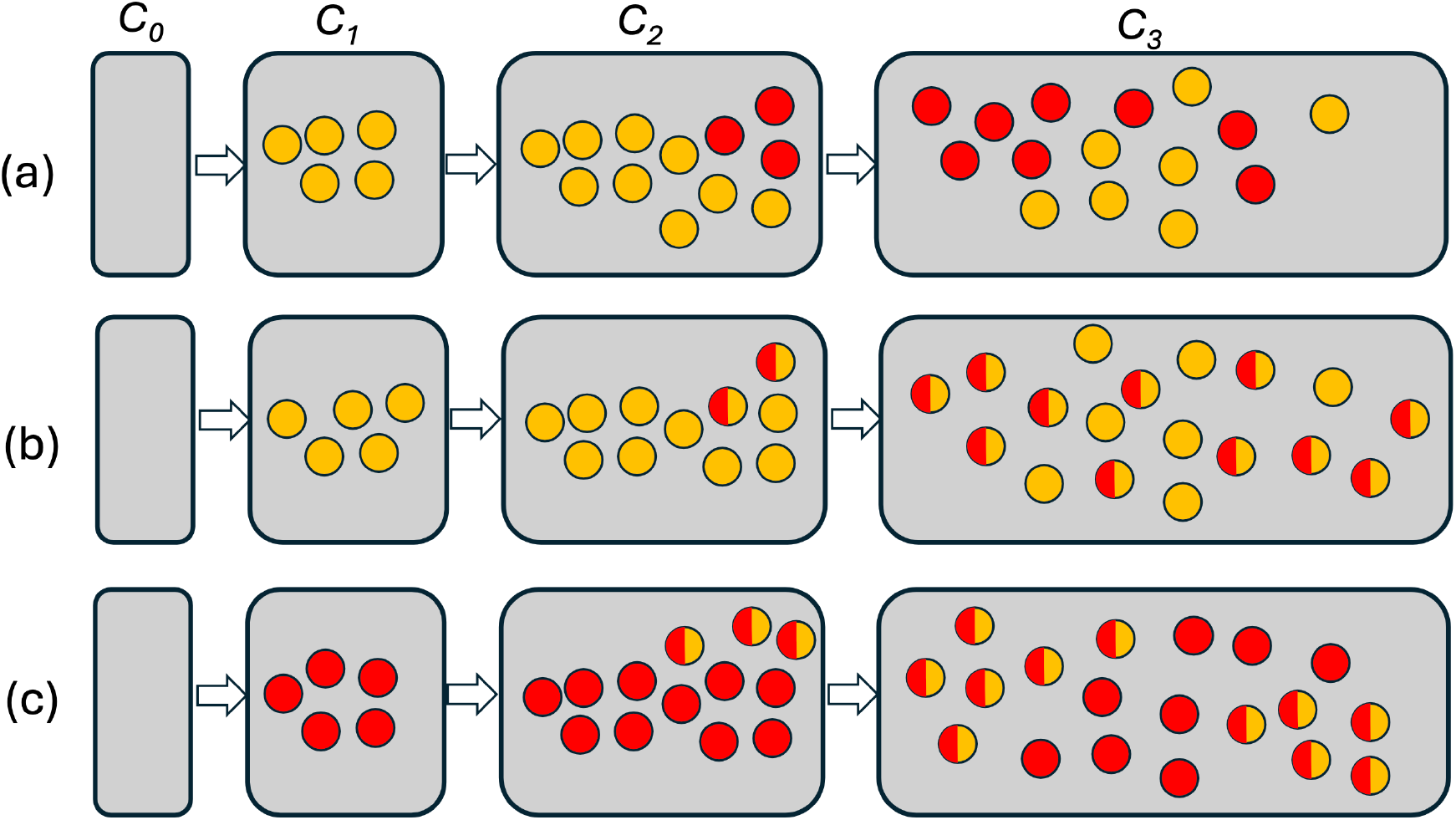
Different applications of the model. The gray boxes represent the compartments LT-HSC, SH-HSC, MPP, etc (*C*_0_, …, *C*_3_) containing the (background) wild-type clone. Circles represent mutants. Yellow is a relatively harmless mutant (TET2), red is a more dangerous mutant (JAK2V617F), and circles containing both colors are double-hit mutants with both mutations. (a) TET2 is introduced in the ST-HSC and JAK2V617F in the MPP compartment; this is an example when the two mutations comprise two different clones. (b) After TET2 is introduced in the MPP compartment and spreads downstream, one of TET2 mutants acquires a secondary JAK2V617F mutation, giving rise to a double-mutant clone. Same as (b) but first a JAK2V617F mutant clone is introduced in the ST-HSC compartment and the secondary TET2 mutations occurs in the MPP compartment.

In what follows we will explore both of these scenarios.

### 3.1 Two consecutive single mutations

Here we consider co-dynamics of two clones in a hematopoietic system, see for example a schematic in figure 4(a). The question that we pose here is whether and how the presence of the harmless mutation may modify the dynamics of the more dangerous mutation.

#### JAK2V617F in the ST-HSC compartment (*C*_1_)

First let us consider the case where the more dangerous, JAK2V617F, mutant clone is introduced in compartment *C*_1_, see figure 5. If no other mutations are present, it will rise from low numbers because *s*^(2)^ *> ŝ*_1_({0, −,− }) (inequality (28)), see panel (a).

**Figure 5:**
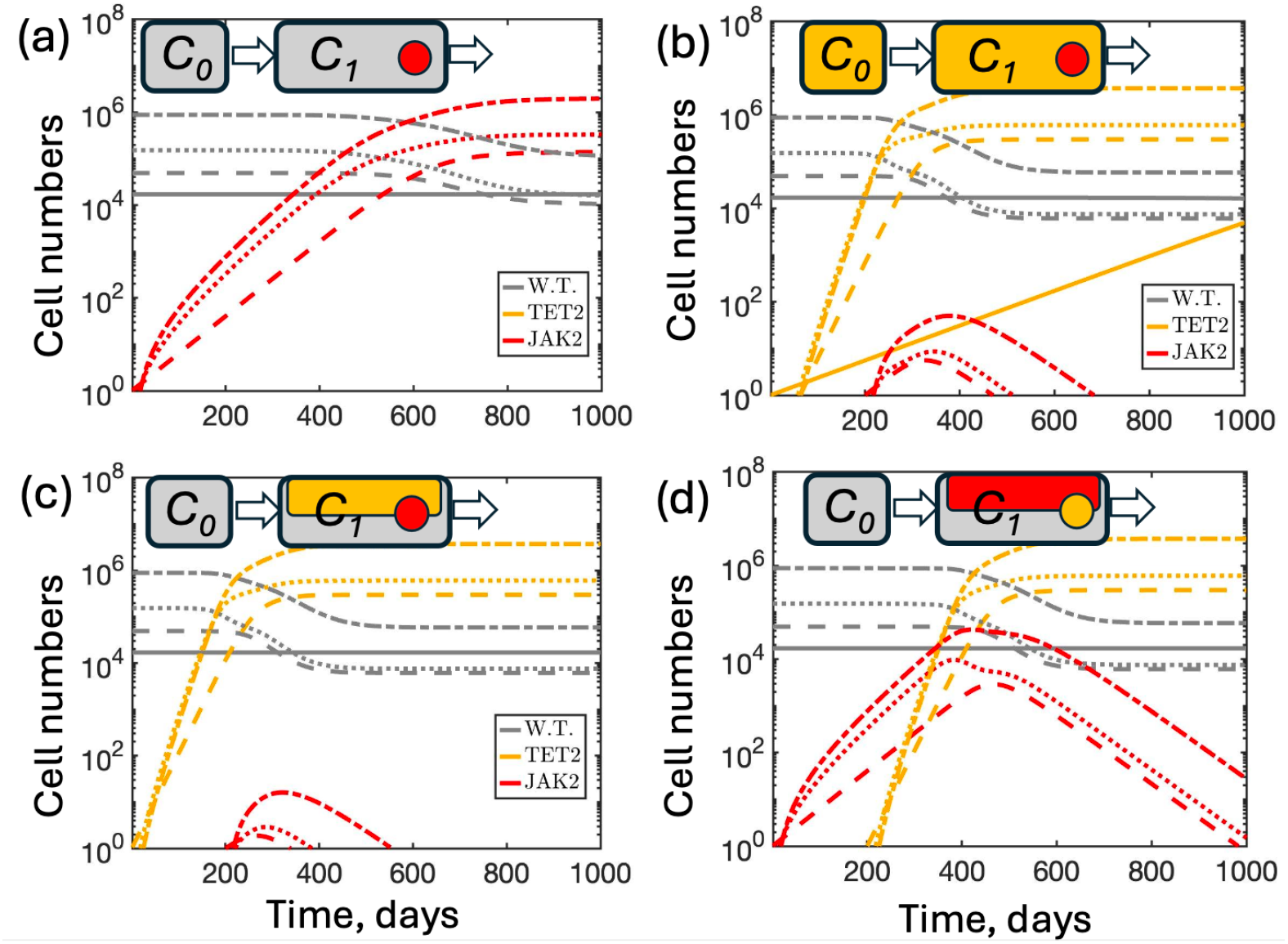
Different scenarios where the JAK2V617F mutant is introduced in the ST-HSC compartment (*C*_1_). The schematics on top of the graphs show the two compartments, LT-HSC and ST-HSC. Established clones are shown by colored rectangles, and a new clone by a circle. Gray color is used for w.t., yellow for TET2, and red for JAK2V617F. The graphs show the abundance of each cell type in each of the compartments (equations (12-14)), with solid, dashed, dotted, and dash-dotted lines corresponding to LT-HSC, ST-HSC, MPP, and CMP/CLP compartments respectively. (a) A single JAK2V617F mutant clone. (b) Before JAK2V617F hits, a TET2 clone is introduced in the LT-HSC compartment, preventing JAK2V617F from rising. (c) Before JAK2V617F hits, a TET2 clone is introduced in the ST-HSC compartment, preventing JAK2V617F from rising. (d) After JAK2V617F hits, a TET2 clone is introduced in the ST-HSC compartment, displacing JAK2V617F. The parameters are given in Table 2, *s*^(*T ET* 2)^ = 0.8 and *s*^(*JAK*2)^ = 0.35. Here and throughout the paper, the simulation horizon is limited to 1000 days, reflecting the approximate maximum physiological lifespan of a laboratory mouse.

Let us next suppose that before the introduction of the JAK2V617F mutant, a mutation of TET2 appeared in the LT-HSC compartment (*C*_0_, the top compartment). Since *s*^(*T ET* 2)^ *>* 0, it will eventually displace the wild type and become the new background clone. The resulting resident state, where TET2 mutants displaced the wild-type, in our notations corresponds to the entry string {−, 0, −, } and the resulting threshold value *ŝ*_1_({−, 0, − }) is given by equation (27). Since it requires a much higher selection coefficient (which exceeds *s*^(1)^ = *s*^(*T ET* 2)^), for our choice of parameters invasion of JAK2V617F in the ST-HSC comparment would not be possible. In figure 5(b) we show a scenario related to this, but not exactly the same. With realistic parameter values, a TET2 mutant with the chosen value *s*^(*T ET* 2)^ cannot fixate in the LT-HSC compartment and reach an equilibrium on time-scales shorter than the mouse life-span (due to the slow rate of LT-HSC division). Therefore, in panel (b) we assumed that a JAK2V617F mutant is generated in the ST-HSC compartment before TET2 has equilibrated. The result however is the same: JAK2V617F fails to expand, just as it would under a TET2 equilibrium.

Two more scenarios are described in figure 5(c,d). In panel (c), a TET2 mutant establishes itself in the ST-HSC compartment, *C*_1_ (because of a higher division rate in the compartment, this can easily happen on a relevant time-scale). Then, a JAK2V617F mutant appears in the same compartment, but it fails to invade because *s*^(*JAK*2)^ *< s*^(*T ET* 2)^ (see invasion requirement (23)). In panel (d), JAK2V617F is introduced in the ST-HSC compartment and invades successfully, but later it is displaced by a TET2 mutant that enters in the same compartment.

#### JAK2V617F in the MPP compartment

Next, let us suppose that the more dangerous mutant, JAK2V617F, is introduced in compartment *C*_2_ (MPP). By assumption, it will expand from low numbers in the wild-type background. If however a TET2 clone is placed in the ST-HSC compartment (*C*_1_) and expands, its presence can change the dynamics of the more dangerous mutant.

The scenario described here (that is, JAK2V617F mutant entering in the MPP compartment), is similar to that described above, where JAK2V617F entered in the ST-HSC compartment, except the calculation of the threshold becomes slightly more cumbersome.

Instead of presenting an explicit formula (such as (27)), we will show the calculation graphically. In figure 3(b) we study outcomes of co-dynamics of two mutants: one entering in the ST-HSC compartment (its fitness, *s*^(1)^, is the horizontal axis of the plot), and the other entering in the MPP compartment (its fitness, *s*^(2)^, is the vertical axis). We explore a range of fitness values for both of these clones. The heat plot presents the equilibrium solution for the clone entering in MPP: above the red line it can invade and get established in MPP (coexisting with the other clone), and below the red line its fitness is below the invasion barrier and it fails to establish a colony. The red line is the invasion barrier for the mutant, such as JAK2V617F, entering in MPP, given that another clone (of fitness *s*^(1)^, e.g. a TET2 mutant) has entered in the SH-HSC compartment.

In our notation, the equilibrium corresponding to the TET2 clone generated in the ST-HSC compartment is given by the entry string {0, 1,− } In order to calculate the threshold value for JAK2V617F in the MPP compartment in the presence of a TET2 clone that entered in the ST-HSC compartment, *ŝ*_2_({0, 1, −}), we first find the solutions 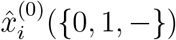 and 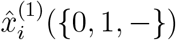, which represent the wild-type and TET2 mutant populations after a “harmless” clone was introduced in the ST-HSC compartment. Then we form 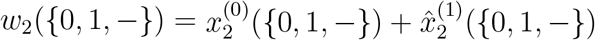 and determine *ŝ*_2_(*{*0, 1, −*}*) = [2*P*_2_(*w*_2_(*{*0, 1, −*}*))]^−1^ − 1.

Figure 3(b) shows that, as *s*^(1)^ increases, the threshold for the new clone in MPP also increases drastically. The sample clonal fitness values used in our illustrative simulations are shown by the dashed white lines: if *s*^(*T ET* 2)^ = 0.8, the value *s*^(*JAK*2)^ = 0.35 is much below the threshold (below the red line) preventing the growth of the JAK2V617F in the MPP compartment in the presence of a TET2 clone that entered in the ST-HSC compartment.

Figure 6(a,b) demonstrates the related dynamics. In panel (a), first a TET2 clone is placed in the ST-HSC compartment and expands, and then a JAK2V617F clone placed in the MPP compartment fails to grow, because *s*^(*JAK*2)^ *< ŝ*_2_({0, 1, − }), see the intersection of the white dashed lines in figure 3(b). In figure 6(b), we start with placing a JAK2V617F clone in the MPP compartment, where it starts growing, until a mutant of TET2 is placed in the ST-HSC compartment, which stunts the growth of JAK2V617F and eventually drives it extinct.

**Figure 6:**
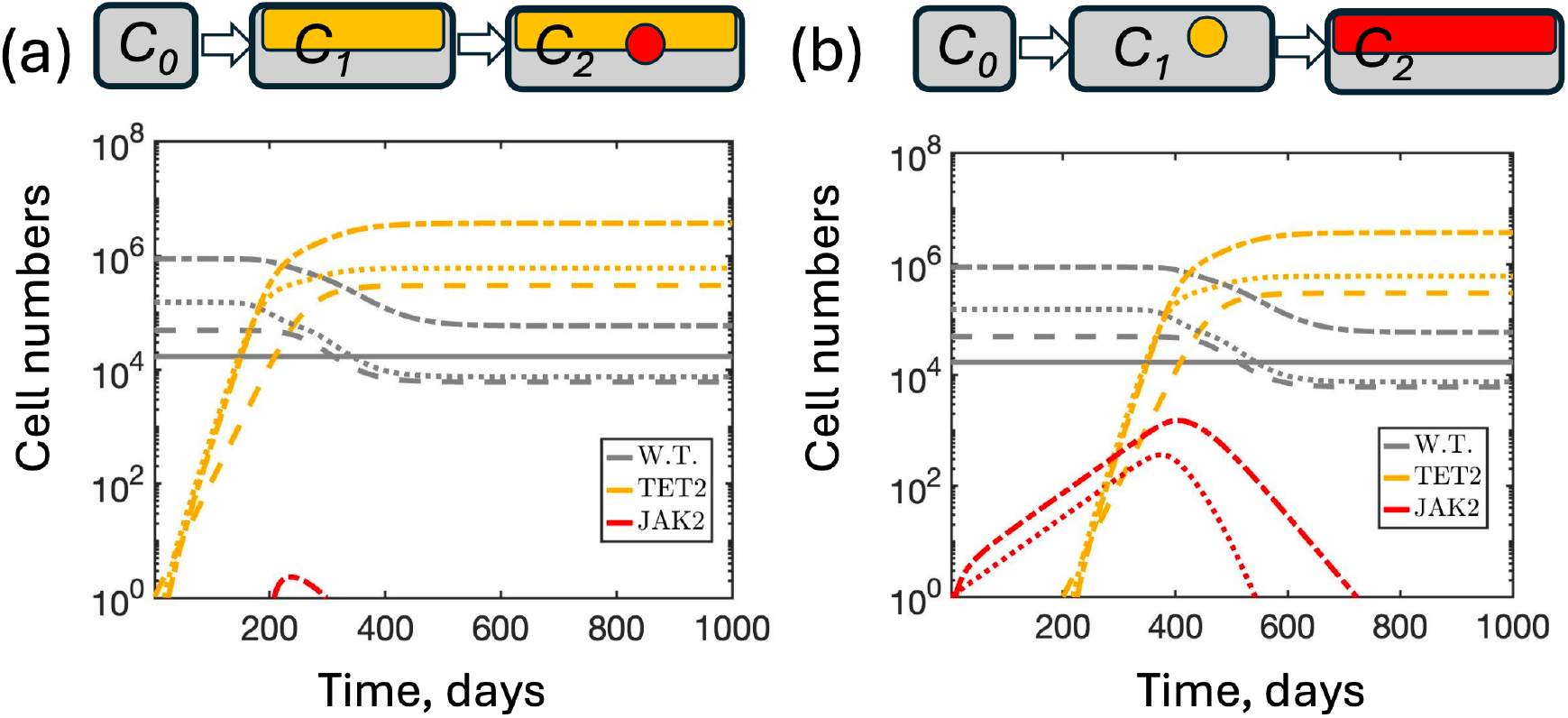
Different scenarios where the JAK2V617F mutant is introduced in the MPP compartment (*C*_2_). (a) Before JAK2V617F hits, a TET2 clone is introduced in the ST-HSC compartment, preventing JAK2V617F from rising. (b) After JAK2V617F hits, a TET2 clone is introduced in the ST-HSC compartment, displacing JAK2V617F. For notations and parameters please see figure 5.

### 3.2 Mutation order in double-mutant evolution

The application below is motivated by experimental studies of patients with myeloproliferative neoplasms (MPNs) carrying both TET2 and JAK2V617F mutations [43]. A central observation was that these mutations frequently coexist within the same hematopoietic clone, indicating that one mutation can arise in a cell that already carries the other. Because the mutations accumulate sequentially, the question naturally arises whether the order in which they are acquired influences disease evolution, see the schematics in figure 4(b,c).

#### Mutation order in JAK2V617F–TET2 myeloproliferative neoplasms: experimental evidence

Studies [42, 43] examined a cohort of human patients with Philadelphia chromosome-negative myeloproliferative neoplasms (MPNs) who carried both a JAK2V617F mutation and a mutation in TET2. The goal was to reconstruct the evolutionary history of the mutant clones in each patient. To this end, hematopoietic progenitor cells were isolated from peripheral blood samples and grown into individual colonies, each originating from a single progenitor cell. Every colony was then genotyped separately for the presence or absence of the two mutations. Since each colony represents the descendants of a single ancestral cell, the collection of colony genotypes provides a snapshot of the clonal lineage within a patient.

The order of mutation acquisition could then be inferred from the observed distribution of colony genotypes. For example, if both TET2-only colonies and TET2/JAK2 double-mutant colonies were present, but no JAK2-only colonies were detected, the most parsimonious evolutionary history is that a TET2 mutation occurred first and that one descendant of the TET2-mutant clone subsequently acquired JAK2V617F. Conversely, the presence of JAK2-only and double-mutant colonies, but not TET2-only colonies, implied the opposite sequence, JAK2 → TET2. Applying this reconstruction procedure across the patient cohort revealed that both mutation orders occur in MPNs, with approximately half of the patients following each evolutionary trajectory.

Having stratified patients according to mutation order, the investigators compared their clinical characteristics. Remarkably, patients in whom JAK2 was acquired first presented with MPN disease more than a decade earlier than patients in whom TET2 was acquired first.

The JAK2-first group also developed a more severe form of the disease, characterized by an increased incidence of polycythemia vera, a higher risk of thrombosis, and greater expansion of blood-cell progenitors, whereas TET2-first patients tended to have a milder clinical course. These observations established that mutation order is not merely a historical record of clonal evolution but an important determinant of disease phenotype. Although TET2 mutations provides the greater intrinsic fitness advantage at the HSC level, the mutational sequence

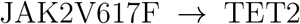

more readily produces aggressive and clinically apparent disease once the second mutation is acquired.

#### Modeling double-hit mutant generation

Our modeling approach can be readily expanded to describe the scenario where double-hit mutants are produced. We will assume that the first clone consists of single-hit mutants (either TET2 or JAK2V617F mutants), and the second clone is generated by one of these cells acquiring a second hit, and consists of double TET2+JAK2V617F mutants.

In figure 7 we assumed additive fitness of the double-hit mutant: if the first and the second hit correspond to mutations that (alone) result in the selection coefficient *s*_1_ and *s*_2_ respectively, the selection coefficient of the double-hit mutant is assumed to be additive:

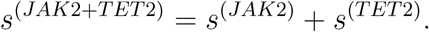

**Figure 7:**
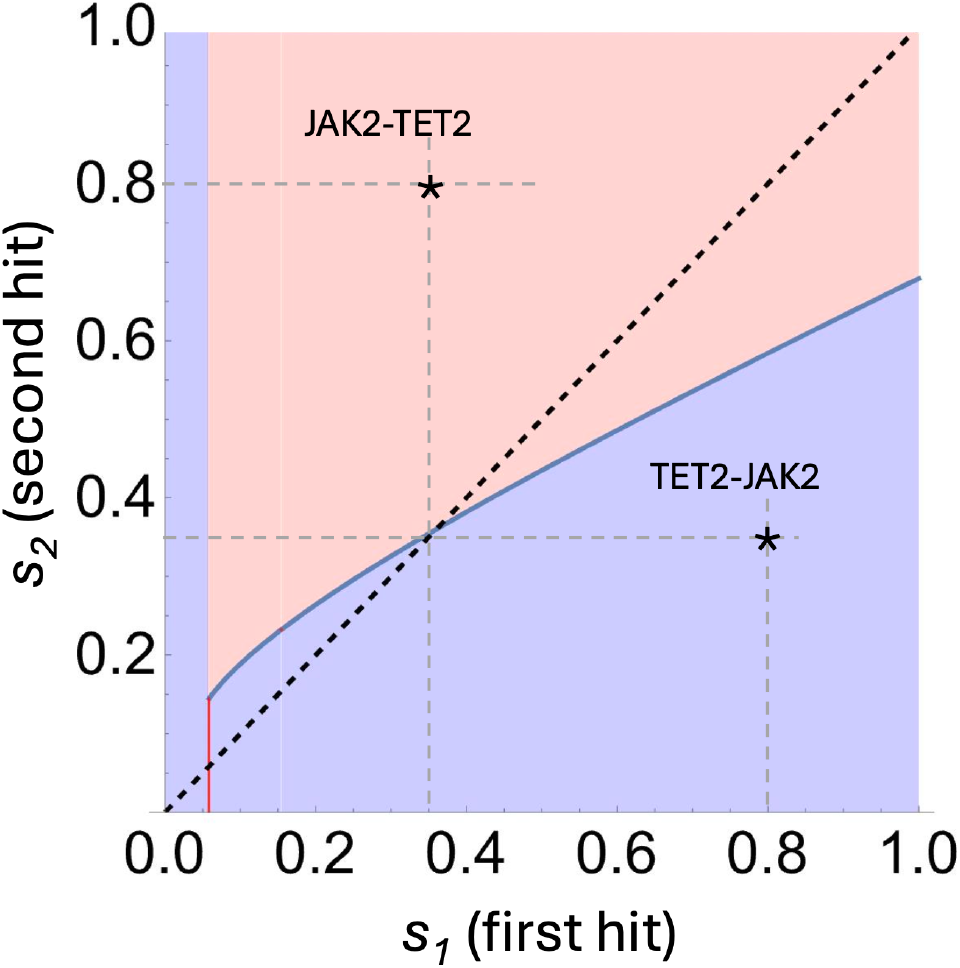
Phase diagram for the invasion of double-hit mutants. Assuming that the first hit arises in the ST-HSC compartment and the second hit in the MPP compartment, the axes are the selection coefficient of the first and the second mutation, while the selection coefficient of the double mutant is assumed to be their sum: the single-hit mutant has fitness *s*^(1)^ = *s*_1_ and the double-hit mutant has *s*^(2)^ = *s*_1_ + *s*_2_. Pink and blue colors correspond to successful invasion and failure of invasion, respectively. The dashed diagonal line shows *s*_1_ = *s*_2_, and the two stars correspond to the two sequences of mutations under the same parameter values as in figure 5.

In our notation, we will study the co-dynamics of two clones. The first clone is a single-hit mutant (either a TET2 or a JAK2V617F mutant). Its fitness is denoted as *s*^(1)^ and is given by the value on the horizontal axis of figure 7. The second clone is the double-hit mutant clone, and its fitness, denoted as *s*^(2)^, is given by *s*^(2)^ = *s*_1_ + *s*_2_. For example, in the case where the first hit is TET2, we have *s*_1_ = *s*^(*T ET* 2)^ and *s*_2_ = *s*^(*JAK*2)^, and the two clones (see figure 4(b)) have fitness *s*^(1)^ = *s*_1_ = *s*^(*T ET* 2)^, *s*^(2)^ = *s*_1_ + *s*_2_ = *s*^(*T ET* 2)^ + *s*^(*JAK*2)^. If the first hit is JAK2V617F, we have *s*_1_ = *s*^(*JAK*2)^ and *s*_2_ = *s*^(*T ET* 2)^, and the two clones (see figure 4(c)) have fitness *s*^(1)^ = *s*_1_ = *s*(*JAK*22), *s*^(2)^ = *s*_1_ + *s*_2_ = *s*^(*T ET* 2)^ + *s*^(*JAK*2)^.

The phase diagram of figure 7 (with the axes *s*_1_ and *s*_2_) shows two regions. The pink region contains parameter combinations that correspond to successful invasion of double-hit mutants, where the double-hit mutant fitness exceeds the invasion threshold imposed by the resident clone. The blue region corresponds to double-hit clone failure, where its selection coefficient is below the threshold.

An example scenario with the selection coefficients of TET2 and JAK2V617F mutants taken 0.8 and 0.35 respectively emphasizes a strong asymmetry with respect to mutational order. In the TET2 then JAK2V617F scenario, the double-hit mutant’s selection coefficient is below the threshold (see the star in the blue region of the diagram). This is because the invasion threshold in this case is very high, due to the high fitness of the first clone, TET2. On the other hand, if the first mutation is JAK2V617F (whose selective coefficient is lower), the invasion threshold it imposes is lower and can be overcome by the double-hit mutant (see the star in the pink region of the diagram).

Figure 8 illustrate the resulting dynamics. In panel (a) we first introduce a TET2 mutant in the ST-HSC compartment (the yellow lines). It spreads through the system, and at time *t* = 200 days we assume that a double-hit mutant is created in the MPP (the yellow and red lines). Since the invasion threshold is very high due to the resident mutant’s high *s*^(1)^ = *s*^(*T ET* 2)^, the double-hit mutant clone’s selection coefficient is below the threshold, and it fails to grow. In panel (b), the order of events is reversed. A JAK2V617F mutant is generated in the ST-HSC compartment and it starts spreading through the system (the red lines), when a second hit (TET2) is acquired in one of the cells with a JAK2V617F mutations in the MPP compartment. The double-hit mutant clone rises because its selection coefficient is above the (comparatively low) invasion threshold. As a result, all three clones (the wild-type, the single-hit JAK2V617F mutants, and the double-hut mutants) coexist in the system.

**Figure 8:**
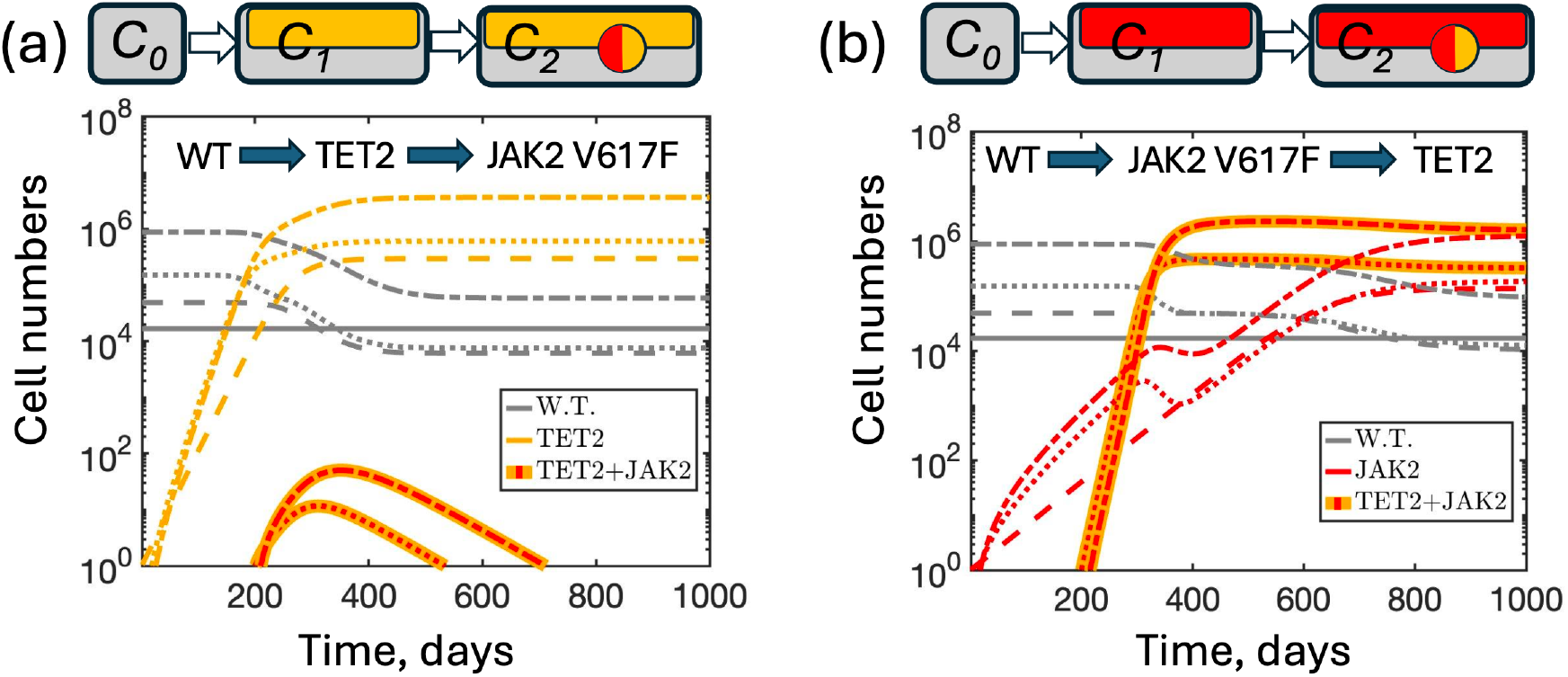
The order of mutations matters. (a) First, a TET2 clone is introduced in the ST-HSC compartment, and then one of its cells in the MPP compartment acquires an additional JAK2V617F mutation, giving rise to a double-hit mutant clone (a thick yellow line with red dashes). This clone cannot invade because its selection coefficient is below the threshold set by the resident TET2 clone. (b) The order of mutations is reversed: first, a JAK2V617F mutation appears in the ST-HSC compartment, and then one of the cells of that clone acquires a TET2 mutation (in MPP). The resulting double-hit mutant clone (a thick red line with yellow dashes) is viable and invades the system, because its selection coefficient is above the (lower) threshold set by the resident JAK2V617F clone. The selection coefficient of the double-hit mutant clone is assumed to be *s*^(1)^ + *s*^(2)^ = *s*^(*T ET* 2)^ + *s*^(*JAK*2)^. For other notations and parameters please see figure 5.

Our numerical experiments should be regarded as a proof of concept rather than as an attempt to represent the process of human MPN clone generation. Recall that the ODE system we are using was parameterized using a mouse model, and in addition, we do not know the exact values of the selection coefficients of the mutants, or how they combine in a double-hit mutant. While in the numerical example of figure 8(a), the double-hit mutant clone with TET2 mutation first fails to grow, this does not mean that it cannot grow in the real system. Its growth might be facilitated by the patient’s aging or inflammation environment, see e.g. [33] where these factors were implemented in a model. The main point our current model makes is that the order of mutations matters because of the different invasion threshold values induced by the first (resident) mutant. Based on this argument, the pathway where JAK2 gene mutates first is expected to result in faster progression of the disease.

##### Remark 1

We note that the above scenario holds under the assumption that the 2nd hit happens in a compartment downstream from where the first mutation occurs, such as ST-HSC for the initial mutation and MPP for the subsequent mutation. If a double-hit mutant happens in the same compartment as the initial (TET2 or JAK2) hit, then the double-hit clone will invade regardless of the order of mutations. This is because the invasion barrier for a clone whose entry compartment coincides with the entry compartment of a previous clone is just the fitness of the previous clone (see expression (22)). In other words, it is enough for the fitness of the new clone to exceed that of the previous clone by any amount. Therefore, no matter what the first mutation is, the double-mutant will have a higher selection coefficient and will out-compete the single mutant in the same entry compartment.

##### Remark 2

If the JAK2V617F → TET2 sequence occurs, the double-hit mutant is more aggressive than the single JAK2V617F mutant, because the JAK2V617F cells in the double-hit mutant clone have a higher self-renewal probability due to the secondary TET2 mutation. If we compare this to the scenario where the two mutations occur in different cells, we have the following interesting result: if generated in a different cell, a TET2 mutation could suppress a JAK2V617F mutant colony, but if it occurs in the same cell, it can promote it.

## 4 Discussion

In this work, we generalized the concept of invasion barriers in a mathematical model of cell evolution in a hierarchically organized tissue, from the case of a single advantageous mutant to populations containing multiple competing clones. Our main finding is that invasion barriers are not fixed properties of the compartments but depend on the composition of the resident population. In particular, every resident clone modifies the effective self-renewal environment experienced by newly arising mutants, thereby changing the conditions under which subsequent mutants can establish themselves. Consequently, the success of a mutant depends not only on its own fitness advantage and the compartment in which it arises, but also on the evolutionary history of the system.

This framework provides a simple mechanistic explanation for history-dependent clonal evolution in hierarchically organized tissues. In our previous work, invasion barriers arose because cells residing in downstream compartments benefit from continuous replenishment from upstream compartments, whereas newly arising mutants do not. Here we showed that once an advantageous clone has become established, it effectively reshapes this ecological landscape for all future mutants. In particular, a highly fit resident clone raises the invasion threshold experienced by later mutants, whereas a resident clone with a smaller fitness advantage creates a lower barrier. Thus, the order in which mutations occur directly influences which evolutionary trajectories remain accessible.

While our analysis can apply to different tissues with hierarchical organization, we parameterized the model specifically for the mouse hematopoietic system due to the availability of detailed kinetic experimental data. Assuming that the general principles of this model also apply to the dyanmnics of hunman hematopoiesis, the mechanisms described here can naturally explain mutation-order effects observed in myeloproliferative neoplasms. Studies have shown that patients carrying both TET2 and JAK2V617F mutations exhibit different clinical phenotypes depending on which mutation occurred first. Our model suggests that this phenomenon does not necessarily require mutation-specific epistatic interactions. Instead, it can emerge from a generic ecological mechanism: the first mutation establishes a resident population that determines the invasion barrier encountered by the second mutation. In the illustrative parameter regime considered here, we assumed that TET2 mutants are characterized by a higher selection coefficient compared to JAK2V617F mutants due to experimental evidence discussed above. In our model, a high-fitness TET2 clone creates a barrier that prevents expansion of a subsequently generated double-mutant clone, whereas a lower-fitness JAK2V617F clone permits invasion after acquisition of a second mutation. The observed asymmetry therefore follows from differences in invasion thresholds rather than from differences in the intrinsic properties of the double-mutant clone itself.

More broadly, the results suggest that clonal evolution in hierarchically organized tissues is fundamentally path-dependent. In classical population genetics, invasion is determined primarily by relative fitness. In contrast, in hierarchical tissues with homeostatic regulation, fitness alone is insufficient to predict evolutionary success. The compartment in which mutations arise, as well as the identity of resident clones and their compartments of entry, together determine whether a newly arising clone can establish itself.

Several simplifying assumptions were made in this study. The model is deterministic and does not explicitly describe stochastic mutation generation or extinction during the earliest stages of clone establishment. Instead, new mutants are introduced as small populations through the initial conditions. In a stochastic setting, the invasion barrier does not delineate “success” from “failure” of a clone, but instead serves as a boundary between supercritical and subcritical populations. We also assumed that mutations act by modifying the effective self-renewal probability and illustrated double-hit mutants using an additive fitness model. In reality, interactions between mutations may be non-additive, and fitness effects may depend on the compartment, as well as age, inflammation, or other features of the microenvironment. In addition, it is important to point out that uncertainties exist regarding the model structure. Model structure can determine which parameters influence the fitness of cell variants. Since in our model, the self-renewal vs differentiation decision is taken during the process of cell division, an increased self-renewal probability renders a cell advantageous, while simply a faster rate of cell division does not. Models in which cell fate decisions are uncoupled from cell division events, or models that include an explicit death rate in stem-cell compartments [13, 44] might also be biologically feasible and can have different properties. While nontrivial invasion barriers in such models are expected to remain, the mathematical expressions would have to be evaluated separately. Similarly, assuming alternative functional forms of the self-renewal regulation (instead of equation (3) used here) will lead to somewhat different functional forms of the invasion barriers. Related to model structure, the exact processes that govern cell fate decisions are uncertain, even for the relatively well-understood hematopoietic system. Due to lack of biological evidence, we assumed that differentiation is uni-directional and that no de-differentiation events occur. If de-differentiation occurs, the relative magnitude of differentiation vs de-differentiation would be an important variable that can influence the results reported here.

Overall, our results identify invasion barriers as a unifying principle governing clonal evolution in hierarchically organized tissues. Rather than being determined by the intrinsic fitness of individual mutations, evolutionary outcomes are shaped by the interaction between hierarchical tissue organization, homeostatic feedback, and historical contingency. This perspective provides a natural explanation for mutation-order effects and suggests that evolutionary history should be regarded as an integral determinant of disease progression.

## Acknowledgments

The authors gratefully acknowledge support of the following grants: DMS 2424853 (DW, NK, TM, AF), and NIH grant R01CA271172 (AF, DW, NK).

